# GeneAutomate: A Browser-Based, Integer-Indexed Platform for Dual-Gene-List Functional Annotation and Interactive Network Visualization

**DOI:** 10.64898/2026.07.16.738882

**Authors:** Rudra Pratap Singh, Amrendra Kumar

## Abstract

Comparative interpretation of two gene lists, for example, two treatment arms, two tissues, or a discovery and a validation cohort, is a routine task in functional genomics. While several tools offer dual-list comparison (e.g., EnrichmentMap, RRHO packages), they typically require local software installation, R/Bioconductor, or manual reconciliation of separate single-list outputs. Most widely used web-based enrichment tools (DAVID, g:Profiler, Enrichr, ShinyGO, WebGestalt) are built around the analysis of a single gene list at a time, and those that support comparison often lack interactive, publication-ready visualization or depend on server-side query latency. Here we present GeneAutomate, a browser-based tool purpose-built for side-by-side comparison of two gene lists. GeneAutomate performs Over-Representation Analysis (ORA) against Gene Ontology (GO) and Reactome using an exact hypergeometric test with Benjamini-Hochberg false discovery rate correction, and Gene Set Enrichment Analysis (GSEA) when ranked (log2 fold-change) input is supplied, alongside Protein-Protein Interaction (PPI) subgraph extraction from BioGRID physical interactions. All reference data (Gene Ontology, Reactome, BioGRID, and NCBI/Ensembl identifier cross-references) are pre-compiled offline into a single integer-indexed database of approximately 32 MB for Homo sapiens, in which every gene identifier Ensembl ID, Entrez ID, official symbol, or alias is resolved to one canonical integer prior to any user query. This design removes live database round-trips from the runtime path, enabling fast, at-your-desk enrichment without installation or a server-side per-query bottleneck. The tool renders thirteen interactive, D3.js- and Cytoscape.js-based comparative visualizations, including a Rank-Rank Hypergeometric Overlap (RRHO) heatmap, a GO-slim “Radar/Spider” functional fingerprint, and chord/edge-bundled cross-talk diagrams that are, to our knowledge, not offered as an integrated set by any existing academic or commercial ORA/GSEA platform. GeneAutomate is an unfunded, individual student project developed with feedback from a professor, and is in its final stage of development. It requires no installation or login. We describe the tool’s architecture, statistical methods, and comparative feature set relative to established academic tools (DAVID, ShinyGO, g:Profiler, Enrichr, WebGestalt, STRING, PANTHER, GeneMANIA, Cytoscape, clusterProfiler, GSEA, Metascape) and commercial platforms (IPA, MetaCore, Pathway Studio, iPathwayGuide, Partek Pathway), and we state candidly the current version’s limitations, which are planned to be the added in next version: single-species (human-only) coverage, no upstream regulator analysis, and comparison currently limited to two (occasionally three) concurrent lists. GeneAutomate is available at https://geneautomate.tech/.

## 1. Introduction

High-throughput transcriptomic, proteomic, and genomic experiments routinely output ranked or unranked gene lists that must be translated into biological hypotheses through functional enrichment analysis. Over-Representation Analysis (ORA), which tests whether a gene list contains more members of a given pathway or GO term than expected by chance, and Gene Set Enrichment Analysis (GSEA), which tests whether a pathway’s genes are non-randomly distributed across a fold-change-ranked list, are the two dominant statistical frameworks in this space [1]. A large ecosystem of tools implements one or both approaches: web servers such as DAVID [2], ShinyGO [3], g:Profiler [4], Enrichr [5], WebGestalt [6], and Metascape [7]; network- and interaction-oriented resources such as STRING [8], GeneMANIA [9], and Cytoscape [10]; ontology- and family-classification resources such as PANTHER [11]; the R/Bioconductor package clusterProfiler [12]; and the original desktop GSEA application distributed by the Broad Institute [1]. In parallel, a distinct tier of commercial platforms QIAGEN Ingenuity Pathway Analysis (IPA), Clarivate MetaCore, Elsevier Pathway Studio, Advaita iPathwayGuide, and Partek Pathway offer curated, often manually reviewed pathway content and additional analytics (e.g., upstream regulator prediction) behind institutional licenses.

Two practical challenges recur across this landscape. First, many established tools are designed around the analysis of a single gene list, making side-by-side comparison of two experimental conditions the most common design in differential expression studies a multi-step, manually intensive process that typically requires running separate analyses and cross-referencing results manually. While several approaches exist for comparative visualization (including multi-list web portals, network-based enrichment mapping, and rank-rank concordance plots), they are often distributed across different platforms, require local software installation, or are not integrated into a single browser-based workflow. Second, enrichment web servers commonly perform each query against a live backend database, which introduces server-side query latency and per-session computational load. GeneAutomate was built to address both gaps directly: (i) a purpose-built dual-gene-list comparative workflow, with visual encodings mirror bar charts, an RRHO heatmap, and cross-talk chord/edge-bundling diagrams specifically designed to show where two lists agree, diverge, or physically interact; and (ii) an architecture in which the entire reference knowledgebase (GO, Reactome, and BioGRID, cross-referenced to a single integer identifier space) is compiled once, offline, into a compact, self-contained database that the analysis backend loads into memory, avoiding live per-query database lookups at runtime. The tool requires no login and runs its full visualization layer as client-side JavaScript in the browser. GeneAutomate is a student-led, unfunded project developed with faculty feedback and is in its final stage of development. It is available at https://geneautomate.tech/.

## 2. Implementation and Architecture

### 2.1 Overview

GeneAutomate is organized into an offline database-compilation layer, a synchronous analysis backend, and a client-side visualization layer. Per the repository’s own language statistics, the codebase is around 80% Javascript and 9% Python reflecting the fact that all thirteen interactive visualizations, and the majority of the interface logic, execute directly in the user’s browser once a compact JSON analysis payload is returned by the backend. The statistical coreidentifier resolution, hypergeometric testing, false-discovery correction, and GSEA running-sum computation is implemented in Python and currently runs as a conventional synchronous web service.

### 2.2 Offline compilation: the integer-core database

The central architectural decision in GeneAutomate is that all biological identifiers are resolved to compact integers during offline compilation, never at query time. This compilation proceeds in three stages. First, a consolidated gene table joining Ensembl IDs, symbols, Entrez IDs, GO annotations, and Reactome pathways is streamed row by row; each novel Ensembl ID is assigned a sequential integer, with the corresponding symbol and Entrez ID registered against the same integer, giving every gene family a single canonical numeric handle. A reverse dictionary (integer→[Ensembl ID, symbol, Entrez ID]) is retained for display. Second, a larger alias tablecontaining previous symbols, alias symbols, and cross-database rescue mappingsis cross-referenced against this integer core. Alias collisions are resolved using an evidence-strength hierarchy: Ensembl, Entrez, and official-symbol mappings are strongest; historical symbols intermediate; alias symbols weakest. When equal-strength sources disagree, the alias is withheld into an “ambiguous” lexicon rather than assigned arbitrarily; the tool reports any user-submitted identifier that falls into this bucket as ambiguous, distinct from unrecognised identifiers. This safeguard ensures that ambiguous mappings fail visibly rather than silently corrupting downstream statistics. Third, the integer core is used to translate GO and Reactome annotation files, the GO DAG, and BioGRID physical interaction records into flat, integer-keyed JSON structures: (a) a GO enrichment matrix mapping each active GO term to its member gene integers; (b) a Reactome enrichment matrix with a single-parent pathway hierarchy (built by taking the first Reactome parent for each pathway and recursively injecting missing ancestor nodes, ensuring no pathway is an orphan in the sunburst hierarchy); (c) an undirected BioGRID PPI adjacency list (excluding non-physical interactions and self-loops); and (d) a GO-slim “biological homes” taxonomy, built by extracting the goslim_generic anchor subset and pre-computing, via breadth-first search over is_a and namespace-matched part_of edges, the nearest slim ancestor(s) for each of the ∼40,000+ GO terms. This slim taxonomy is used exclusively for the GO-slim functional fingerprint visualisation (Section 3) and is kept separate from the full-resolution GO term matrix used for ORA statistics, so enrichment p-values are always computed against the complete GO term universe rather than the coarser slim categories.

The compiled assetsthe integer map, GO and Reactome matrices and lexicons, PPI adjacency list, and GO-slim taxonomyform the tool’s reference database, approximately 32 MB for Homo sapiens. All files are minified (compact JSON, no whitespace) at compilation. The backend loads this database once at process startup into memory-resident Python dictionaries and sets, so every subsequent analysis request performs only in-memory set intersections and integer array lookups, with no file or network I/O.

The 32 MB size results from four compounding design choices. First, the integer-core identifier scheme replaces every long-form gene identifier with a compact integer wherever it appears in GO, Reactome, and PPI matrices; because a given gene recurs across thousands of term-to-gene associations, this is the largest single contributor to size reduction. Second, all compiled JSON assets are written with strict minification removing whitespace and indentation. Third, the pipeline merges redundant annotation tablesgene symbol, Ensembl, Entrez cross-references, GO annotations, and Reactome annotationsinto a single unified gene table before matrix construction, so identifier translation happens once per gene rather than being re-derived or restored inside each downstream matrix. Fourth, the compiled database is loaded from disk with a single json.load() call at backend startup and held in memory for the lifetime of the process; every subsequent request is served from these in-memory structures, so no per-request disk I/O or database connection occurs. This in-memory model is only practical because the database is compact enougha multi-gigabyte reference compiled without these steps would not fit in memory on typical server hardware.

### 2.3 Runtime analysis pipeline

At query time, a user-submitted gene list (Ensembl IDs, Entrez IDs, official symbols, or common aliasesoptionally paired with a log2 fold-change value per gene) is resolved against the integer core using a three-tier lookup: (i) the merged normalization map, returning a canonical integer on success; (ii) the ambiguous lexicon, if the identifier was flagged as unresolvable during compilation; or (iii) an “unmapped” bucket if the identifier is not recognized at all.

For each list, GeneAutomate then computes ORA against both the GO and Reactome enrichment matrices using an exact hypergeometric test, evaluated in log-space via the log-gamma function to avoid numerical overflow on large gene universes, followed by Benjamini-Hochberg false discovery rate (FDR) correction [13] applied independently within the GO and Reactome term families. Term-size gating (configurable minimum overlap size, and minimum/maximum term size) is applied before correction to avoid testing under-powered or overly generic terms. The background gene universe defaults to the full annotated gene set for each ontology, but a user-supplied custom background list is also supported, in which case both the query and the annotation universe are restricted to the intersection with that background before testinga feature offered by some but not all comparable ORA tools. When per-gene log2 fold-change values are supplied, GeneAutomate additionally computes a GSEA-style running-sum enrichment statistic for each list’s top-ranked significant terms: genes are ranked by fold-change, and a running score is incremented by a fixed step for each ranked gene present in the term’s gene set and decremented by a complementary step otherwise, producing an enrichment curve and the rank positions (“barcode”) of hits along it. In its current implementation, this running-sum statistic is unweighted by fold-change magnitude (i.e., it corresponds to the classical, magnitude-independent Kolmogorov-Smirnov-type running sum rather than a weighted variant), a detail we note explicitly here for methodological transparency. To keep browser-side rendering responsive, each enrichment curve is downsampled to a bounded number of points using a peak-preserving algorithm that always retains local maxima/minima and the curve endpoints.

Results from the two (or more) input lists are then merged into shared GO and Reactome term maps, keyed by term ID, capturing each list’s q-value, overlap size, and overlapping genes, and ranked by the best (minimum) q-value across lists; the top-ranked 150 GO terms and 150 Reactome pathways are retained for downstream visualization to bound response payload size. The union of genes appearing in these top-ranked terms is then used to extract a PPI subgraph: for every pair of genes in this union, an edge is drawn if a physical BioGRID interaction exists between them, and each gene is annotated with its degree centrality within this subgraph, a precomputed PageRank score (where available), and whether it belongs to the top 5% most-connected “hub” genes of the full compiled interactome (a cutoff fixed at compilation time). Each gene is also tagged with a bitmask recording which of the input lists it belongs to, enabling the bar, radar, chord, and network visualizations to consistently color-code genes by list membership from a single shared data structure.

**Figure 1.**
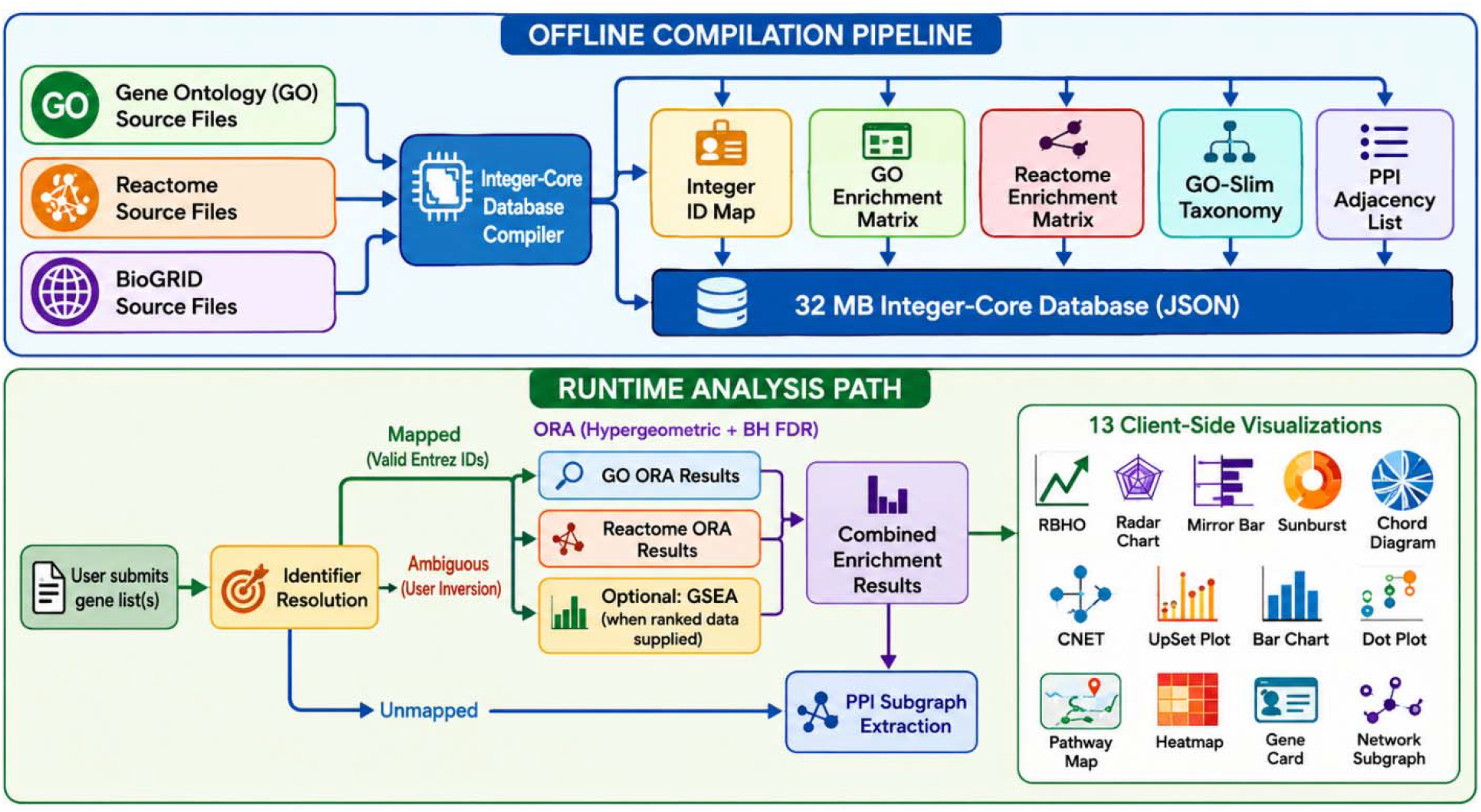
Schematic overview of the GeneAutomate workflow.

## 3. Results and Key Features

### 3.1 A dual-list-first analytical design

Unlike single-list-first tools that require re-running the entire workflow to compare two conditions, GeneAutomate’s backend natively processes an arbitrary-keyed payload of gene lists in a single request, computing per-list statistics and then fusing them into shared term maps, a shared gene ledger, and a shared PPI subgraph. The interface is designed and tested around comparing two gene lists side by side; a proportional Venn/set-overlap diagram of significantly enriched GO terms is additionally supported for up to three concurrent lists, beyond which the tool explicitly declines to render an area-proportional Venn diagram (since such diagrams are not geometrically well-defined beyond three sets) and instead informs the user that higher-order intersection support (an UpSet plot) is planned for a future version, rather than attempting a misleading or degenerate rendering.

### 3.2 Thirteen interactive, comparative visualizations

All visualizations are rendered client-side using D3.js (for statistical/geometric charts) and Cytoscape.js (for force-directed and circular network layouts), and support interactive hover, zoom, and click-to-isolate behavior. GeneAutomate currently ships thirteen distinct visualization techniques across five analysis panels:

1. **Proportional Venn Overlap**. Area-proportional overlap of significantly enriched GO terms across up to three input lists, with an explicit fallback message (rather than a misleading render) when more lists are supplied.
2. **Diverging Mirror Bar Chart**. A back-to-back bar chart contrasting the top enriched terms of each list, allowing at-a-glance comparison of which pathways dominate in list A versus list B.
3. **RRHO Heatmap**. A Rank-Rank Hypergeometric Overlap plot [14] comparing the two lists as continuous fold-change-ranked profiles rather than fixed-threshold gene sets, revealing threshold-independent concordance or discordance between the two conditions.
4. **CNET / Gene-Concept Network**. A bipartite network linking enriched terms to their member genes, with genes color-coded by list membership.
5. **Enrichment Map Network**. A term-term similarity network (nodes = enriched terms, edges = shared-gene overlap), in the tradition of the Cytoscape EnrichmentMap approach [15], rendered natively in-browser.
6. **Semantic Treemap**. A space-filling hierarchical treemap of the same enriched-term set, offered as an alternate, area-encoded view toggled from the same panel as the Enrichment Map network.
7. **Hierarchical Sunburst Partition**. A radial partition of enriched Reactome pathways following Reactome’s own pathway hierarchy, with each Reactome pathway visualization enforcing single-parent tree topology so that every active pathway resolves to an unbroken path back to the ontology root.
8. **Sector Pathway Profile (Polar Lollipop)**. A polar lollipop chart summarizing pathway-level enrichment strength within each hierarchical sector.
9. **Cross-Talk Network (Hierarchical Edge Bundling)**. An edge-bundled radial network highlighting Reactome pathways that are jointly implicated by both gene lists.
10. **Hub Gene Chart**. A ranked bar chart of the highest-centrality (hub) genes within the extracted PPI subgraph.
11. **Force-Directed Network (fCoSE)**. An interactive, physically simulated layout of the full PPI subgraph, with Markov Clustering (MCL) used to isolate functionally coherent network modules.
12. **Cross-Talk Analysis (circular cross-talk diagram)**. A chord-diagram-style circular layout isolating direct physical PPI edges that bridge list-A-specific and list-B-specific genes, filtering out intra-list edges to foreground cross-talk between the two conditions.
13. **GO-Slim Radar / Spider Fingerprint**. A radial (spider) plot of each list’s functional composition across GO-slim “biological home” categories, allowing the overall functional character of two gene lists to be compared as a single visual silhouette rather than a term-by-term list.

To our knowledge, the RRHO heatmap, the GO-slim Radar/Spider Fingerprint, and the dual-list PPI cross-talk chord-style diagram are not offered together, as an integrated set, in any other free or commercial ORA/GSEA web tool surveyed here; RRHO is otherwise available as a standalone R/Bioconductor package [14] rather than as part of an integrated enrichment web platform, and GO-slim composition is more commonly presented as a stacked bar chart than as a radial fingerprint.

**Figure 2.**
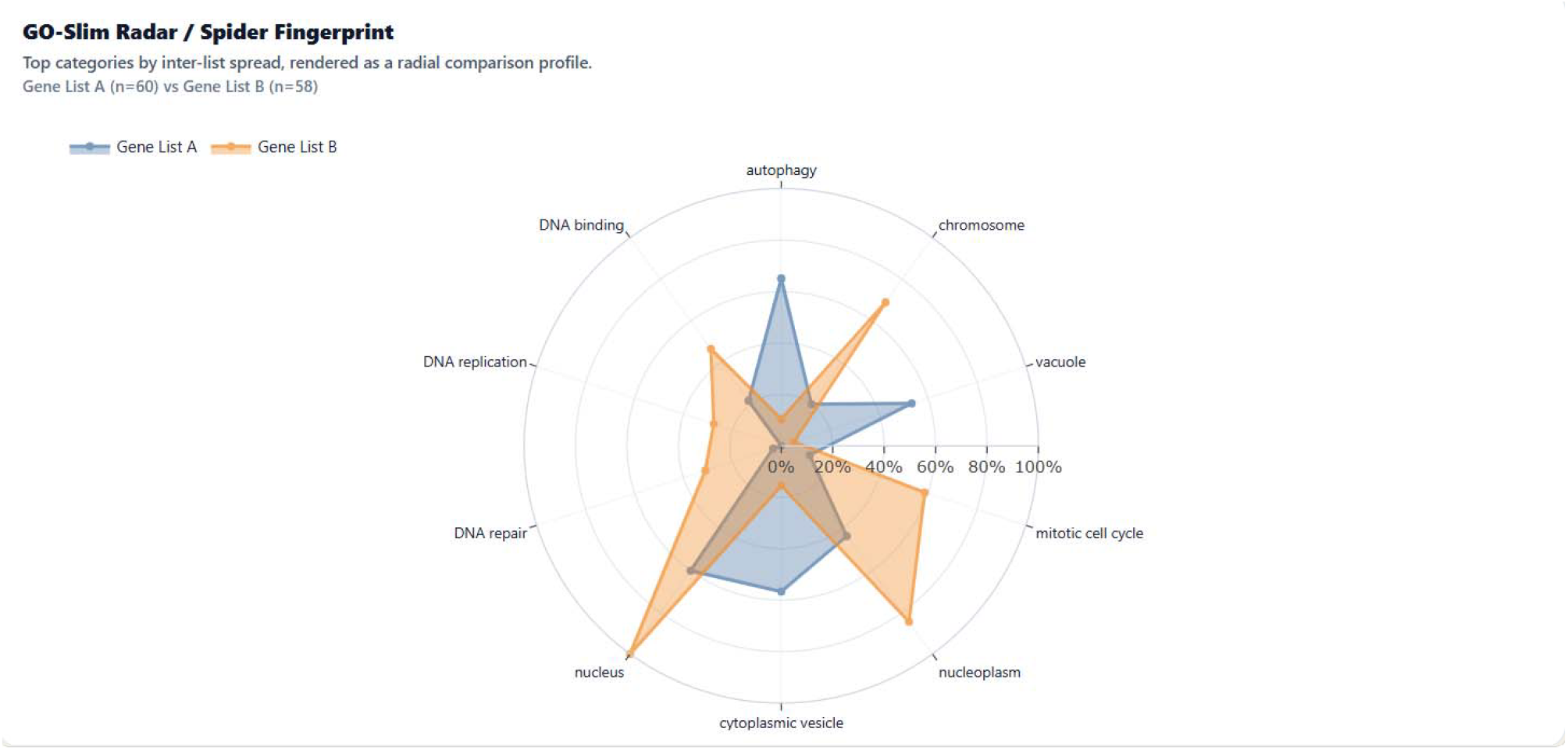
GO-Slim Radar / Spider Fingerprint. Radial and interactive comparison of the functional (GO-slim) composition of two input gene lists, with each axis representing one GO-slim “biological home” category and each list rendered as a distinct overlapping polygon.

**Figure 3.**
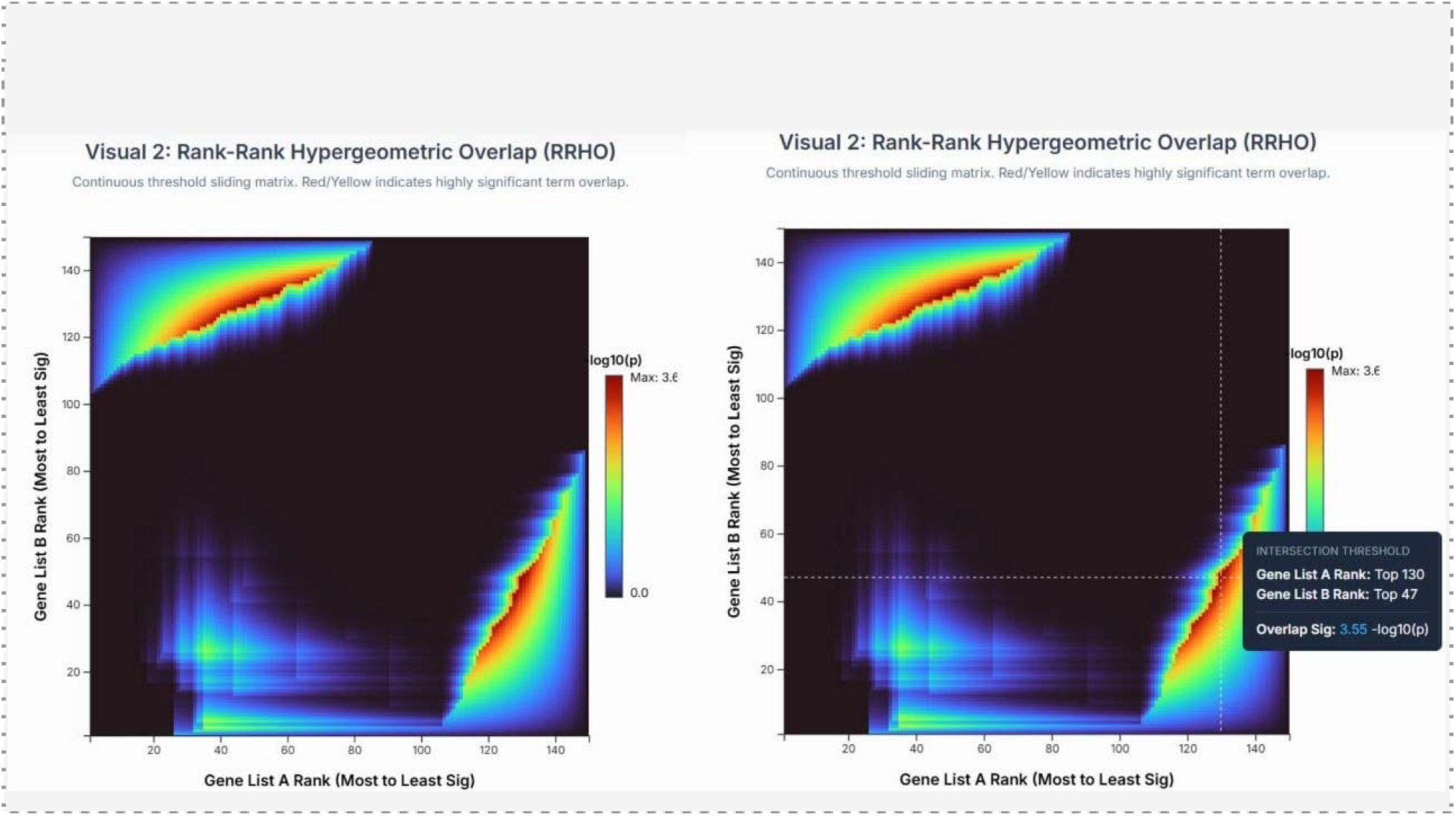
Rank-Rank Hypergeometric Overlap (RRHO) heatmap comparing two fold-change-ranked gene lists as continuous profiles, illustrating threshold-independent regions of concordant and discordant enrichment.

**Figure 4.**
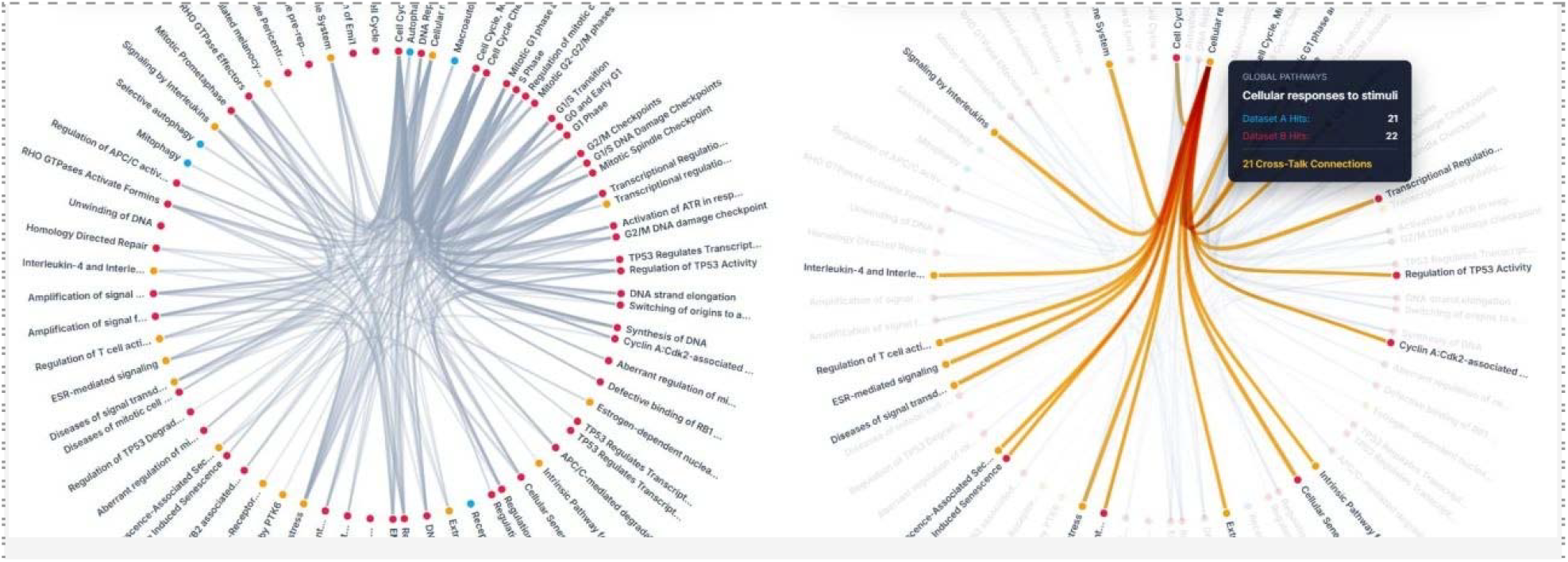
Cross-Talk Network (Hierarchical Edge Bundling) view of Reactome pathways jointly implicated across the two input gene lists.

**Figure 5.**
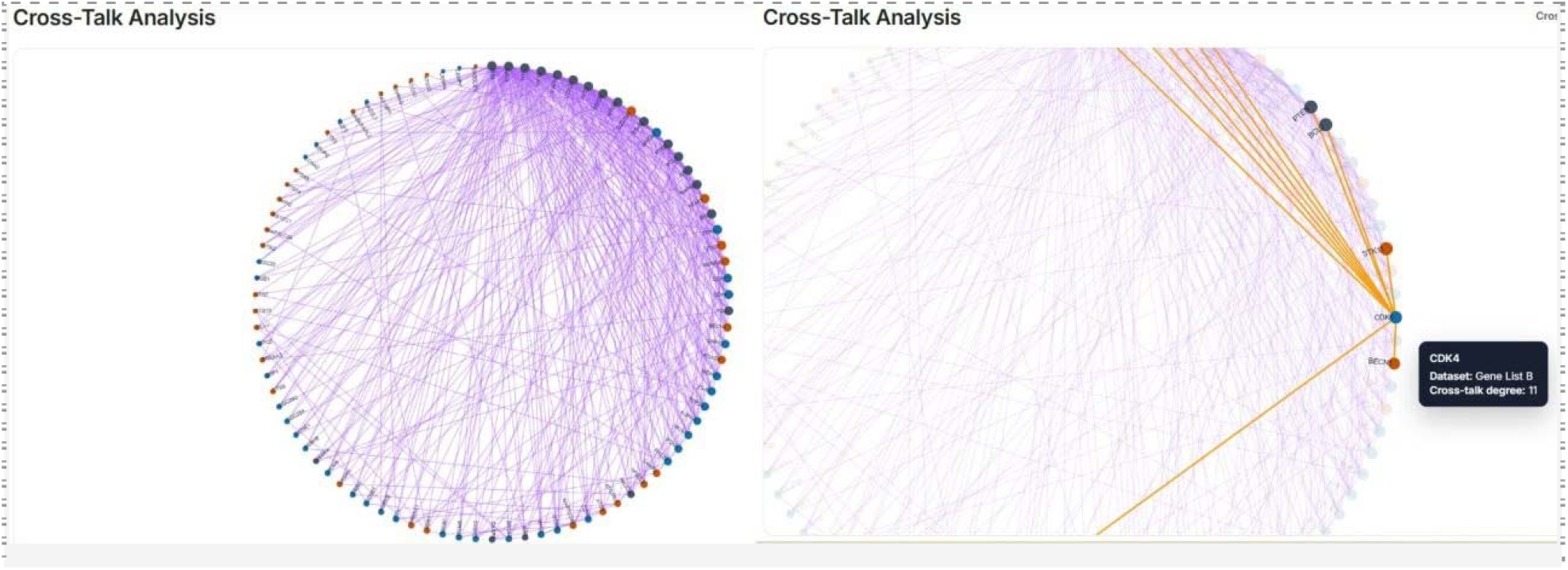
Cross-Talk Chord-style diagram of the extracted BioGRID physical protein-protein interaction subgraph, isolating direct interactions that bridge genes unique to each of the two input lists.

### 3.3 Statistical rigor and transparency

GeneAutomate computes an exact hypergeometric p-value (not a normal or chi-square approximation) for every tested term, using log-gamma arithmetic to remain numerically stable on large gene universes, and applies the standard Benjamini-Hochberg step-up FDR procedure [13] with proper monotonicity enforcement. Every returned enrichment result carries its overlap size (k), term size (M), and both raw p-value and corrected q-value, and every analysis additionally reports which input identifiers were successfully mapped, which were flagged as ambiguous (multiple candidate genes with no way to disambiguate from the evidence available), and which were entirely unrecognizeda level of identifier-resolution transparency that is not uniformly surfaced by comparable web tools.

### 3.4 User convenience and cost

GeneAutomate requires no installation, no account registration or login, and no local software (R, Java, or a Cytoscape desktop installation) to access its full comparative visualization suite; a user can paste two gene lists and receive interactive comparative results within a single browser session.

## 4. Discussion

GeneAutomate occupies a specific niche within the crowded functional-enrichment tool landscape: it is, to our knowledge, one of a small number of openly accessible web tools that treat multi-gene-list comparison as a first-class workflow rather than a secondary feature bolted onto a single-list-first designa category that also includes Metascape [7], which offers zero-installation multi-list upload and cross-list heatmap/network summaries. Within that category, GeneAutomate differentiates itself primarily through its comparison-specific visual encodingsthe RRHO heatmap, the GO-slim radar/spider fingerprint, and the PPI cross-talk diagramswhich are not, to our knowledge, offered by Metascape or by other zero-installation multi-list web portals surveyed here. Its integer-core, offline-compiled reference database is intended to reduce runtime dependency on live external database queries, and its client-side visualization layer, built on D3.js and Cytoscape.js, provides a level of interactivity (hover, zoom, click-to-isolate) that static-image-output tools do not offer. The tool’s thirteen visualization types, several of which (RRHO, GO-slim radar fingerprint, PPI cross-talk diagrams) are uncommon in combination elsewhere, are intended to make dual-condition comparison visually native rather than an afterthought requiring manual reconciliation of two separate single-list reports.

## 5. Methods

### 5.1 Data sources

GeneAutomate’s compiled reference database for Homo sapiens integrates the following primary data sources, each retrieved at the version current as of the dates indicated:

- Gene Ontology core directed acyclic graph (OBO-format release), providing GO term definitions, namespaces, and the is_a / part_of / regulates relationship edges used for GO-slim ancestor resolution.
- GO Consortium generic GO-slim term subset, used as the anchor set for the GO-slim “biological homes” taxonomy underlying the Radar/Spider Fingerprint visualization.
- UniProt-GOA human gene-association file (GAF format), providing gene-to-GO-term annotations; rows carrying a “NOT” qualifier (i.e., annotations that explicitly exclude a gene from a term) are filtered out during compilation.
- NCBI Entrez Gene gene2go annotation file, contributing to the consolidated gene-annotation table.
- Reactome Ensembl2Reactome and NCBI2Reactome identifier-mapping files, together with the Reactome pathway hierarchy (parent-child relation file) and human pathway name listings, used to build the Reactome enrichment matrix and single-parent pathway hierarchy.
- BioGRID protein and genetic interaction dataset, filtered at compilation time to physical-interaction records only, used to build the PPI adjacency list underlying the interactome and cross-talk visualizations.

These source files were filtered, cross-referenced against one another, and merged offline into a small number of consolidated intermediate tables (a unified gene annotation table and a filtered BioGRID physical-interaction table), which were in turn compiled into the final integer-indexed JSON assets described in Section 2.2. The complete compiled database totals approximately 32 MB for the human reference set. As all source files were the most current releases available at the time of compilation (May 2026), we treat this date as the effective version reference for the underlying annotation content.

### 5.2 Statistical methods

Over-representation testing uses the hypergeometric distribution, with the p-value for observing k or more overlapping genes between a query set of size n and a term of size M, drawn from a background population of size N, computed exactly via log-gamma-based binomial coefficients and summed over the upper tail. Multiple-testing correction uses the Benjamini-Hochberg procedure [13], applied separately within the GO and Reactome term families, with q-values enforced to be monotonically non-decreasing as required by the step-up procedure. Terms are excluded from testing if their background size falls outside a configurable minimum/maximum term-size window, and results are excluded from reporting if the observed overlap falls below a configurable minimum overlap count; both thresholds default to conservative values (minimum overlap = 3 genes; term-size window = 5–1000 genes) but can be relaxed by the user, and a custom background gene universe may be supplied in place of the whole-annotation background. When ranked (log2 fold-change) data are supplied, a GSEA-style running-sum statistic is computed for each list against its own top-ranked significant terms, using equal, unweighted step sizes for hits and misses (the classical, magnitude-independent variant of the running-sum statistic), producing both the full enrichment curve and the rank positions of contributing genes.

### 5.3 Software implementation

The offline database-compilation pipeline and the runtime analysis backend are implemented in Python. The client-facing applicationincluding all thirteen visualizationsis implemented in JavaScript using D3.js for statistical and geometric chart types and Cytoscape.js for force-directed and circular network layouts, with HTML/CSS for layout and styling; per the repository’s own reported language composition, this JavaScript/HTML/CSS layer constitutes approximately 92% of the codebase by volume. The tool is deployed as a public web application and does not require any client-side installation.

## 6. Availability and Future Directions

GeneAutomate is freely accessible at https://geneautomate.tech, with no registration or login required to run an analysis. The source code, including the database-compilation scripts described in Section 2, will be made available via the website’s About page and will be fully open-sourced upon release of a stable version; in the interim, the code is available from the corresponding author upon reasonable request.

Planned future development includes: (i) extension beyond Homo sapiens to additional model organisms; (ii) implementation of upstream transcriptional regulator analysis; (iii) full multi-list (greater than two) comparative visualization across all chart types, building on the backend’s existing bitmask-based multi-list data model and complementing the current three-list Venn/set-overlap ceiling with UpSet-style high-order intersection plots; and (iv) a systematic, blinded benchmarking study of runtime performance and statistical concordance against established ORA/GSEA tools. We welcome community feedback via the corresponding author to help prioritize this roadmap.

## References

1. Subramanian A, Tamayo P, Mootha VK, Mukherjee S, Ebert BL, Gillette MA, Paulovich A, Pomeroy SL, Golub TR, Lander ES, Mesirov JP. Gene set enrichment analysis: a knowledge-based approach for interpreting genome-wide expression profiles. Proc Natl Acad Sci USA. 2005;102(43):15545–15550.

2. Sherman BT, Hao M, Qiu J, Jiao X, Baseler MW, Lane HC, Imamichi T, Chang W. DAVID: a web server for functional enrichment analysis and functional annotation of gene lists (2021 update). Nucleic Acids Res. 2022;50(W1):W216–W221.

3. Ge SX, Jung D, Yao R. ShinyGO: a graphical gene-set enrichment tool for animals and plants. Bioinformatics. 2020;36(8):2628–2629.

4. Kolberg L, Raudvere U, Kuzmin I, Adler P, Vilo J, Peterson H. g:Profiler—interoperable web service for functional enrichment analysis and gene identifier mapping (2023 update). Nucleic Acids Res. 2023;51(W1):W207–W212.

5. Kuleshov MV, Jones MR, Rouillard AD, Fernandez NF, Duan Q, Wang Z, Koplev S, Jenkins SL, Jagodnik KM, Lachmann A, et al. Enrichr: a comprehensive gene set enrichment analysis web server 2016 update. Nucleic Acids Res. 2016;44(W1):W90–W97.

6. Wang J, Vasaikar S, Shi Z, Zhang B, Greer M. WebGestalt 2017: a more comprehensive, powerful, flexible and interactive gene set enrichment analysis toolkit. Nucleic Acids Res. 2017;45(W1):W130–W137.

7. Zhou Y, Zhou B, Pache L, Chang M, Khodabakhshi AH, Tanaseichuk O, Benner C, Chanda SK. Metascape provides a biologist-oriented resource for the analysis of systems-level datasets. Nat Commun. 2019;10:1523.

8. Szklarczyk D, Kirsch R, Koutrouli M, et al. The STRING database in 2023: protein-protein association networks and functional enrichment analyses for any sequenced genome of interest. Nucleic Acids Res. 2023;51(D1):D638–D646.

9. Warde-Farley D, Donaldson SL, Comes O, Zuberi K, Badrawi R, Chao P, Franz M, Grouios C, Kazi F, Lopes CT, et al. The GeneMANIA prediction server: biological network integration for gene prioritization and predicting gene function. Nucleic Acids Res. 2010;38(suppl_2):W214–W220.

10. Shannon P, Markiel A, Ozier O, Baliga NS, Wang JT, Ramage D, Amin N, Schwikowski B, Ideker T. Cytoscape: a software environment for integrated models of biomolecular interaction networks. Genome Res. 2003;13(11):2498–2504.

11. Mi H, Ebert D, Muruganujan A, Mills C, Albou LP, Mushayamaha T, Thomas PD. PANTHER version 16: a revised family classification, tree-based classification tool, enhancer regions and extensive API. Nucleic Acids Res. 2021;49(D1):D394–D403.

12. Wu T, Hu E, Xu S, Chen M, Guo P, Dai Z, Feng T, Zhou L, Tang W, Zhan L, et al. clusterProfiler 4.0: a universal enrichment tool for interpreting omics data. Innovation (Camb). 2021;2(3):100141.

13. Benjamini Y, Hochberg Y. Controlling the false discovery rate: a practical and powerful approach to multiple testing. J R Stat Soc Series B. 1995;57(1):289–300.

14. Plaisier SB, Taschereau R, Wong JA, Graeber TG. Rank-rank hypergeometric overlap: identification of statistically significant overlap between gene-expression signatures. Nucleic Acids Res. 2010;38(17):e169.

15. Merico D, Isserlin R, Stueker O, Emili A, Bader GD. Enrichment map: a network-based method for gene-set enrichment visualization and interpretation. PLoS One. 2010;5(11):e13984.

16. Ashburner M, Ball CA, Blake JA, Botstein D, Butler H, Cherry JM, Davis AP, Dolinski K, Dwight SS, Eppig JT, et al. Gene ontology: tool for the unification of biology. Nat Genet. 2000;25(1):25–29.

17. Gene Ontology Consortium. The Gene Ontology knowledgebase in 2023. Genetics. 2023;224(1):iyad031.

18. Gillespie M, Jassal B, Stephan R, Milacic M, Rothfels K, Senff-Ribeiro A, Griss J, Sevilla C, Matthews L, Gong C, et al. The Reactome pathway knowledgebase 2022. Nucleic Acids Res. 2022;50(D1):D687–D692.

19. Oughtred R, Rust J, Chang C, Breitkreutz BJ, Stark C, Willems A, Boucher L, Leung G, Kolas N, Zhang F, et al. The BioGRID database: a comprehensive biomedical resource of curated protein, genetic, and chemical interactions. Protein Sci. 2021;30(1):187–200.

20. QIAGEN Ingenuity Pathway Analysis (IPA). QIAGEN Digital Insights. Available at: https://digitalinsights.qiagen.com/products-overview/discovery-insights-portfolio/analysis-and-visualization/qiagen-ipa/

21. Clarivate MetaCore. Clarivate. Available at: https://clarivate.com/

22. Elsevier Pathway Studio. Elsevier. Available at: https://www.elsevier.com/

23. Advaita iPathwayGuide. Advaita Bioinformatics. Available at: https://www.advaitabio.com/

24. Partek Pathway. Partek Incorporated. Available at: https://www.partek.com/

